# A genetically encoded, pH-sensitive mTFP1 biosensor for probing lysosomal pH

**DOI:** 10.1101/2020.11.04.368654

**Authors:** Marcus Y. Chin, Anand R. Patwardhan, Kean-Hooi Ang, Austin L. Wang, Carolina Alquezar, Mackenzie Welch, Phi T. Nguyen, Michael Grabe, Anna V. Molofsky, Michelle R. Arkin, Aimee W. Kao

## Abstract

Lysosomes are important sites for macromolecular degradation, defined by an acidic lumenal pH of ∼4.5. To better understand lysosomal pH, we designed a novel, genetically encoded, fluorescent protein (FP) based pH biosensor called FIRE-pHLy (<u>F</u>luorescence Indicator <u>RE</u>porting <u>pH</u> in <u>Ly</u>sosomes). This biosensor was targeted to lysosomes with lysosomal-associated membrane protein 1 (LAMP1) and reported lumenal pH between 3.5 and 6.0 with monomeric teal fluorescent protein 1 (mTFP1), a bright cyan pH sensitive FP variant with a pKa of 4.3. Ratiometric quantification was enabled with cytosolically oriented mCherry using high-content quantitative imaging. We expressed FIRE-pHLy in several cellular models and quantified the alkalinizing response to bafilomycin A1, a specific V-ATPase inhibitor. In summary, we have engineered FIRE-pHLy, a specific, robust and versatile lysosomal pH biosensor that has broad applications for investigating pH dynamics in aging and lysosome-related diseases, as well as in lysosome-based drug discovery.

---

Lysosomes support diverse cellular functions by acting as sites of macromolecular degradation, nutrient recycling, pathogen clearance and signaling events that regulate cellular functions ^1–4^. Mammalian cells eliminate misfolded proteins using either the ubiquitin-proteasome system or autophagy-lysosome pathway. Both play indispensable roles in protein quantity and quality control in the cell ^5,6^. The degradative abilities of lysosomes are conferred by an acidic lumen (pH ∼4.5-4.7) ^7,8^ which contains more than fifty hydrolytic enzymes, also known as “acid hydrolases” that break down major macromolecules into building blocks that are recycled for cellular reuse ^9–11^. Lysosomal acidity is maintained through the vacuolar-type H^+^-ATPase (V-ATPase) proton pump, an evolutionarily conserved electrogenic pump that generates a proton gradient across membranes by coupling proton translocation with ATP hydrolysis ^12^. Additional contributions to lysosomal pH setpoint are made by a number of counter-ion channels and transporters ^13^.

Lysosomal pH dynamics are broadly implicated in biological and disease pathways. Loss-of-function mammalian V-ATPase mutations are embryonic lethal ^14^, highlighting the significance of lysosomal function, in particular pH, to the sustainment of life. In cancer, aberrant V-ATPase activity is linked to hyper-acidic lysosomes that promote tumor proliferation and invasion ^15–17^. Even relatively small alterations in the proton concentration (∼0.5 – 0.9 pH units) can have dramatic effects on tumor aggressiveness ^18,19^. In contrast, loss of lysosomal acidity is observed in aging. Yeast vacuoles (metazoan homolog of lysosomes) and *C. elegans* lysosomes lose their acidity with increasing age ^20–22^, but can be rescued with caloric restriction that upregulates V-ATPase activity ^21^. Additionally, neuronal health is highly regulated by lysosomal function, as demonstrated by insights from human genetics that link lysosomal dysfunction to a wide range of neurological diseases ^23,24^. Notably, reduced lysosomal pH is a probable key factor in the pathogenesis of familial forms of Alzheimer’s disease, Parkinson’s disease, prion diseases and amyotrophic lateral sclerosis ^25–28^. Furthermore, Alzheimer’s disease-related presenilin-1 mutations have been shown to prevent proper acidification of lysosomes by inhibiting assembly of V-ATPase subunits ^28–30^. These studies highlight the importance of investigating lysosomal pH regulatory mechanisms in disease. Collectively, these findings have transformed our understanding of lysosomes from passive waste receptacles to dynamic participants in regulating cellular health and disease, thus making them salient therapeutic targets ^_31_^.

Given the central role of pH in lysosomal function and overall cellular homeostasis, numerous types of lysosomal probes have been developed. Several small-molecule pH-sensitive dyes, organic fluorophores and synthetic probes (e.g. LysoSensor, LysoTracker, FITC-dextran, pHrodo-dextran, DAMP, quantum dots) label and measure lysosomal pH within cells ^8,28,32–36^. Wolfe *et al*., 2013, compared the most frequently used pH probes for their sensitivity, localization and reported the limitations encountered for accurately quantifying the very low pH values of lysosomes. However, these probes have disadvantages due to their poor specificity of subcellular targeting and cytotoxicity (e.g. LysoSensor Yellow/Blue DND-160 function at shorter wavelengths, excitation-329nm/emission-440nm) that lead to autofluorescence and imaging artifacts, modification of cellular metabolic activity, and leakage from cells ^28,32,37,38^.

On the other hand, genetically encoded pH biosensors based on fluorescent proteins (FP) have many advantages such as (i) controlled expression in different cell types and tissues, (ii) enhanced intracellular specificity and (iii) bypassing of dye-incubation steps to (iv) enable long-term, live imaging studies in cells and animals. The first genetically encoded intracellular pH biosensors (called ‘pHluorins’) were developed through directed mutations of specific residues of green fluorescent protein (GFP) to pH-sensitive histidine residues ^39^. The chromophores of FP are sensitive to protons revealing correlations between pH and fluorescent readout ^40^. Genetically encoded biosensors have emerged as essential tools for probing cellular ions including Ca^++ 41^, H^+ 39^, Zn^2+ 42^, Cl^- 43^, Mg^2+ 44^, and K^+ 45^. Several pH-sensitive FPs have been described and targeted to inaccessible environments such as organelle lumens to measure the pH of various intracellular compartments within the secretory-endocytic pathway. Previously characterized biosensors include EGFP (pKa 6.0) to map endosomal acidification ^46^, pHRed (pKa 6.6) to measure intracellular pH ^38^, pHuji (pKa 7.7) for imaging exo- and endocytosis ^47^, and Keima (pKa 7.7) ^48^, GFP-LC3 (pKa 6) or mRFP-LC3 (pKa 4.5) for detection of autophagy ^49^. Additionally, Burgstaller *et al*. utilized the cyan FP variant mTurquoise2 (pKa = 3.1) to develop a Förster resonance energy transfer (FRET) based biosensor to measure pH throughout the endomembrane system ^50^.

Recently, two ratiometric biosensors targeted to lysosomes using lysosomal-associated membrane protein 1 (LAMP1) have been published with the following expression cassettes: (i) mCherry-pHluorin-mouseLAMP1 ^51^ and (ii) sfGFP-ratLAMP1-mCherry fusions ^52^. Both biosensors used LAMP1 for lysosomal targeting, but different topologies of FPs for pH sensing. The described probes have a reported pKa of ∼6.5 and ∼5.9, respectively. Topologically, the Ponsford *et al*. probe positioned both FP domains within the lysosome lumen while in the design of Webb *et al*. the pH-sensing sfGFP and the mCherry domain face the lumen and cytosol, respectively. Because the physiological pH of the lysosome is ∼4.5, a sensor with a more acidic pKa could be more suitable for reporting the acidic pH range of lysosomes for wide-range applications.

Using the diverse toolkit of FPs ^53,54^, we engineered a mTFP1-humanLAMP1-mCherry construct, which is a dual-fluorescent cyan/red fusion protein that is targeted to lysosomes to report lysosomal pH. We call this biosensor **<u>F</u>**luorescence **<u>I</u>**ndicator **<u>RE</u>**porting **<u>pH</u>** in **<u>Ly</u>**sosomes, or ‘**FIRE-pHLy**’ ^55^. FIRE-pHLy showed specificity with respect to lysosomal localization and for measuring pH within a range of 3.5 to 6.0, with a calculated pKa of 4.4. The biosensor responded to lysosome alkalinizing agents and demonstrated a dynamic pH response in a variety of cell types. High-content imaging of FIRE-pHLy allowed us to measure thousands of cells per condition and precisely quantify these responses. Given the emerging attention to lysosomal pH in neurodegeneration and aging, we explored the utility of FIRE-pHLy in the context of primary neurons, human induced pluripotent stem cells (iPSCs), and neuroblastoma cells. To our knowledge, FIRE-pHLy is the first lysosome-targeted pH biosensor that incorporates mTFP1 as its pH-sensing domain, allowing for pH measurements within the highly acidic range of physiological lysosomes. FIRE-pHLy was adapted to *in vitro* cellular models using both traditional imaging and high-content analysis.

## RESULTS AND DISCUSSION

### Design principles for a ratiometric lysosomal pH biosensor

To develop a reliable lysosomal pH biosensor, we have selected a ratiometric system in which the relative brightness of two reporters is used to quantify pH measurements. In this type of ratiometric, dual-reporter system, one fluorophore changes its signal in response to proton concentration while the other serves as a stable reference point for identifying lysosomes and normalizing fluorescent signals. This capability represents a significant advantage over single fluorophore biosensors that can lead to biased measurements between samples or experiments ^56,57^.

For our purposes, a ratiometric lysosomal pH reporter required the following features: (1) a domain for lysosomal targeting, (2) a cytosolically facing fluorescent protein that exhibits stable brightness at physiological intracellular pH (pH range 6.8 to 7.2) ^7^ and (3) a lysosomal lumen-facing fluorescent protein that provides dynamic lysosomal pH sensing at highly acidic pH (pH <5.0). For lysosomal targeting, we utilized LAMP1, a type 1 membrane protein harboring a tyrosine-based lysosomal sorting motif in its short cytoplasmic tail (last 5 amino acids ‘GYQTI’) ^58,59^. For the cytosolic, pH-insensitive domain of the reporter, we tested a number of candidates and ultimately chose mCherry for its brightness and fluorescent stability at physiological intracellular pH ranges and is described in previous ratiometric studies ^56,60–62^.

The success of a lysosomal pH biosensor depends upon identifying a fluorescent protein that accurately reflects the highly acidic pH of the lysosome. The ideal fluorescent protein for this purpose required a low pKa to allow for pH sensing within the anticipated lysosomal pH range from ∼3.5 to 6.0. Additional major attributes in choosing a pH-sensitive fluorescent protein include high brightness, photostability and the ability to maintain proper protein folding and integrity within the acidic lysosomal environment. After testing different candidates, we selected mTFP1 (monomeric teal fluorescent protein 1). A variant of cyan fluorescent protein, mTFP1 possesses a pKa of 4.3 as well as a robust sigmoidal pH response, as measured in cell-free conditions, across a broad acidic and alkaline pH range ^63^. Additionally, mTFP1 resists common FP pitfalls such as photobleaching and aggregation ^64^. Thus, mTFP1 offers a suitable balance of favorable attributes for the pH-sensitive aspect of a ratiometric pH biosensor. The physicochemical properties of mTFP1 and mCherry are described in Table 1 ^62,63,65^.

**Table 1.**
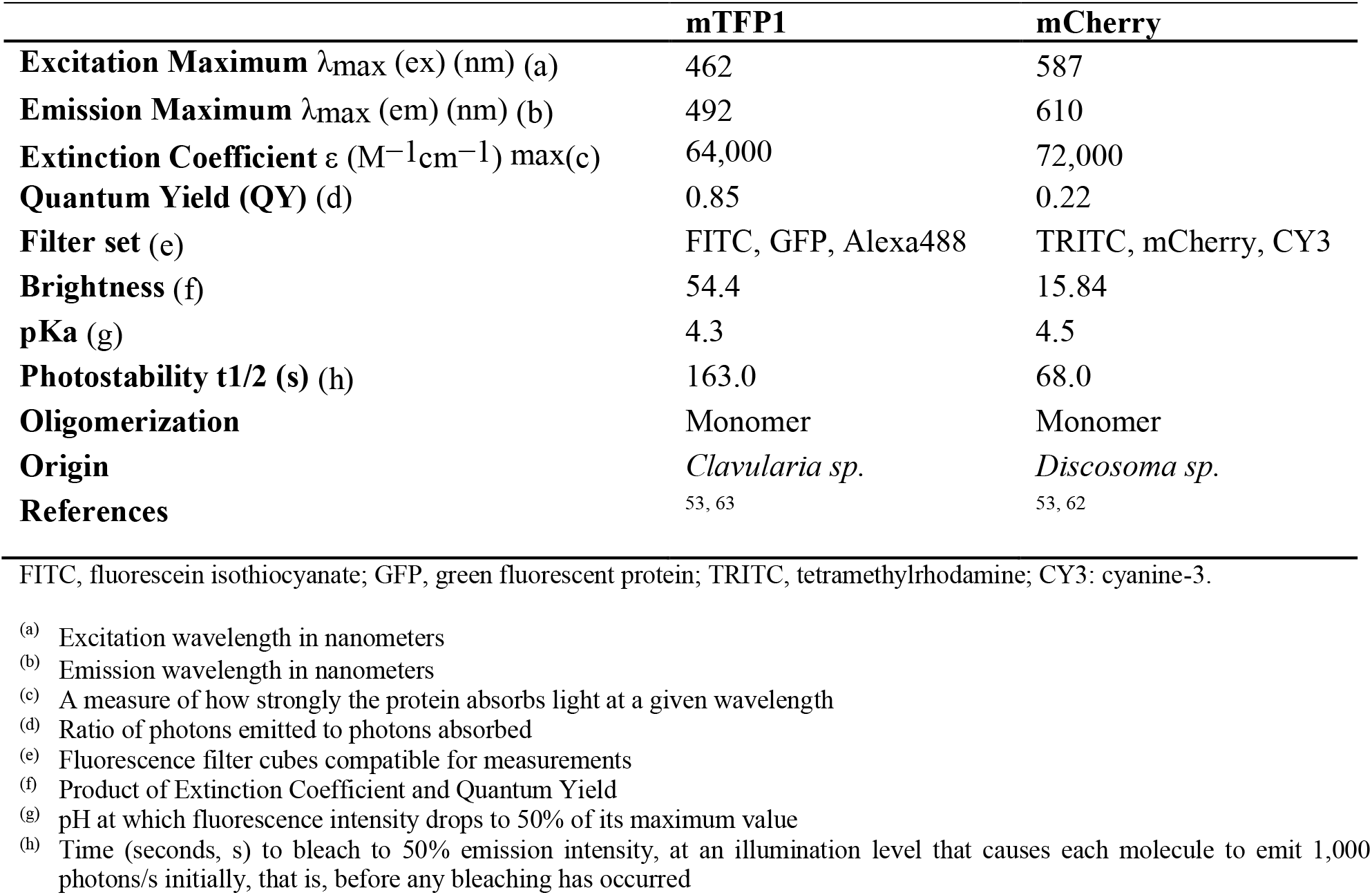
Physicochemical properties of mTFP1 and mCherry.

|  | **mTFP1** | **mCherry** |
| --- | --- | --- |
| **Excitation Maximum** $\lambda_{\text{max}}$ (ex) (nm) (a) | 462 | 587 |
| **Emission Maximum** $\lambda_{\text{max}}$ (em) (nm) (b) | 492 | 610 |
| **Extinction Coefficient** $\epsilon$ ($\text{M}^{-1}\text{cm}^{-1}$) max(c) | 64,000 | 72,000 |
| **Quantum Yield (QY)** (d) | 0.85 | 0.22 |
| **Filter set** (e) | FITC, GFP, Alexa488 | TRITC, mCherry, CY3 |
| **Brightness** (f) | 54.4 | 15.84 |
| **pKa** (g) | 4.3 | 4.5 |
| **Photostability** t1/2 (s) (h) | 163.0 | 68.0 |
| **Oligomerization** | Monomer | Monomer |
| **Origin** | *Clavularia sp.* | *Discosoma sp.* |
| **References** | 53, 63 | 53, 62 |

FITC, fluorescein isothiocyanate; GFP, green fluorescent protein; TRITC, tetramethylrhodamine; CY3: cyanine-3.
(a) Excitation wavelength in nanometers
(b) Emission wavelength in nanometers
(c) A measure of how strongly the protein absorbs light at a given wavelength
(d) Ratio of photons emitted to photons absorbed
(e) Fluorescence filter cubes compatible for measurements
(f) Product of Extinction Coefficient and Quantum Yield
(g) pH at which fluorescence intensity drops to 50% of its maximum value
(h) Time (seconds, s) to bleach to 50% emission intensity, at an illumination level that causes each molecule to emit 1,000 photons/s initially, that is, before any bleaching has occurred

The assembled chimeric fluorescent protein construct consisted of an N-terminal, lysosomal lumen-facing, pH-sensitive mTFP1 fused to the transmembrane portion of human LAMP1 (hLAMP1) and a C-terminal, pH-insensitive mCherry outside the lysosome (**Fig. 1A,1B**). A flexible linker (GGSGGGSGSGGGSG), rich in small and polar amino acids, was added between mTFP1 and LAMP1 to promote correct protein folding and retention of biological and fluorescence properties ^66^. To allow correct sorting, maintain a fixed distance between the two proteins and minimize mCherry aggregation, a rigid linker (PAPAPAP) was placed between LAMP1 and mCherry ^66,67^. Expression of the construct was driven by the human cytomegalovirus (CMV) or human ubiquitin C (UbC) promoter cloned within pLJM1 lentivirus backbone. We designated the resulting chimeric fluorescent protein as FIRE-pHLy, for <u>F</u>luorescence Indicator <u>RE</u>porting <u>pH</u> in <u>Ly</u>sosomes.

**Figure 1.**
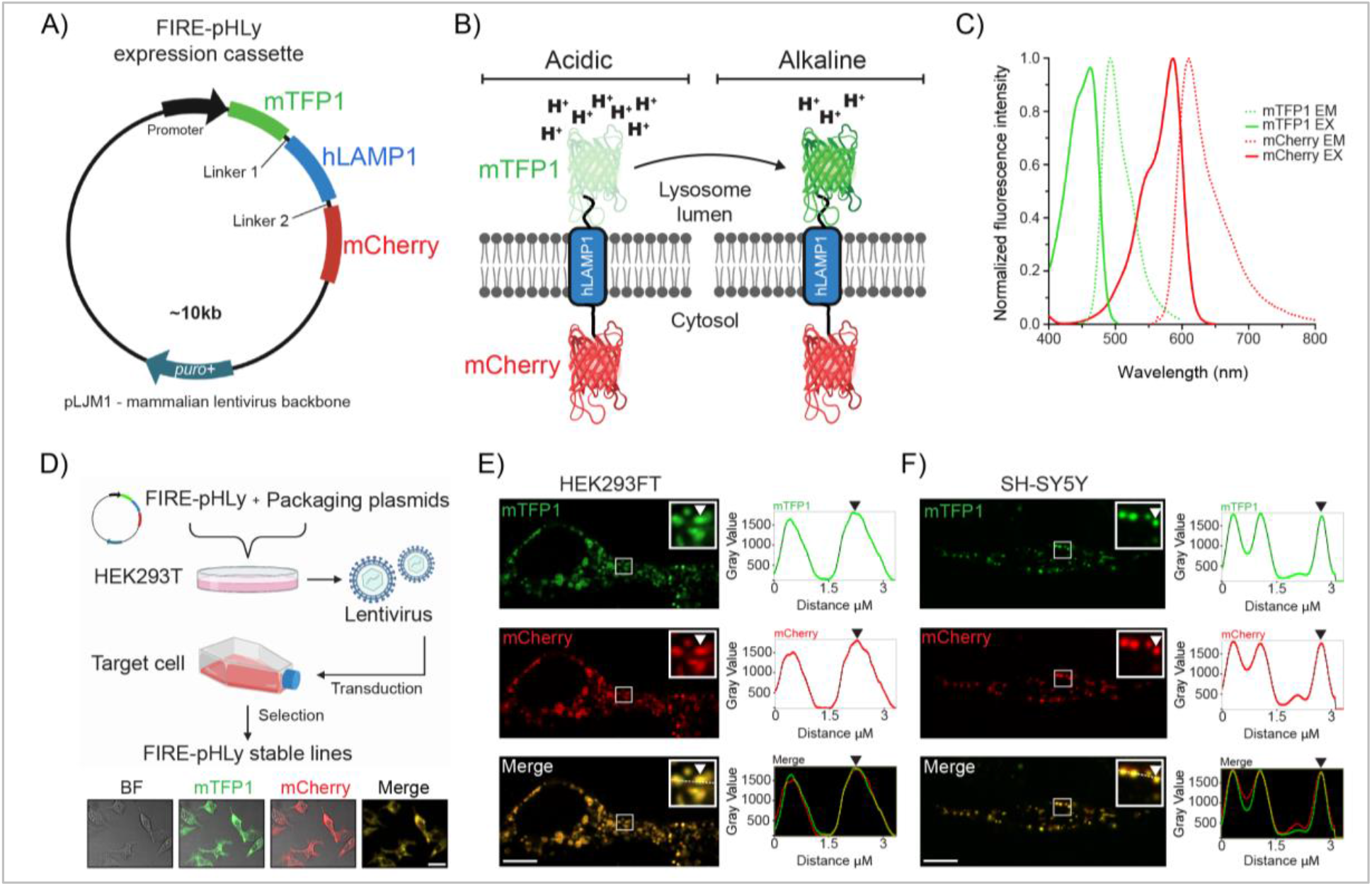
Design of FIRE-pHLy, a ratiometric lysosomal pH biosensor. **(A)** Design of FIRE-pHLy expression cassette driven by the CMV promoter (in HEK293FT cells) or human UbC promoter (in SH-SY5Y cells) cloned in the lentiviral pJLM puromycin-resistant plasmid. Chimeric protein (N- to C-terminus) mTFP1-hLAMP1-mCherry is targeted to lysosomes via the type-I transmembrane human LAMP1 peptide sequence. Linker regions 1 (GGSGGGSGSGGGSG) and 2 (PAPAPAP) allow proper folding and expression of each protein portion. **(B)** Representation of FIRE-pHLy expressed on lysosomal membranes and mTFP1 fluorescence levels in acidic and alkaline conditions. Lysosomal pH-sensitive mTFP1 located within the lumen and lysosomal pH-insensitive mCherry is located on the cytosolic side. **(C)** Excitation (solid lines) and emission spectra (dashed lines) for mTFP1 and mCherry. The 470 nm and 587 nm laser lines were used to excite mTFP1 and mCherry, respectively. Spectra values were obtained and adapted from FPbase ^53^. Refer to Table 1 for physicochemical properties of FIRE-pHLy FPs. **(D)** Workflow of generating stable FIRE-pHLy cell lines using lentiviral vectors. Representative low magnification confocal fluorescence images of brightfield (BF), mTFP1 (green), mCherry (red) and merged channels (yellow) in stable FIRE-pHLy-expressing HEK293FT cells. Scale bar = 25 μm **(E-F)** Live imaging frames of FIRE-pHLy expressing stable cells **(E)** HEK293FT and **(F)** SH-SY5Y with zoomed inset highlighting mTFP1 and mCherry puncta (white arrowhead) and corresponding linescan intensity profile measured along the white line (right panel). Scale bars = 10 μm.

Spectral compatibility is important in dual-color, ratiometric reporters. **Fig. 1C** shows the reported peak excitation and emission wavelengths for mTFP1 (462 and 492 nm, respectively) and mCherry (587 and 610 nm, respectively) ^53,62,63^. To assess bleed-through, we experimentally compared the crosstalk and cross-excited mTFP1 and mCherry with both 470 nm and 587 nm laser lines. mTFP1 was excited at 470 nm and detected in the mCherry channel. Similarly, mCherry was excited at 587 nm and detected in the mTFP1 (green) channel (**Fig. S1**). In both the cases, the results show minimal crosstalk, demonstrating that mTFP1 and mCherry exhibited suitable spectral compatibility for ratiometric imaging.

Using lentiviral transduction, FIRE-pHLy was stably expressed in human embryonic kidney 293 (HEK293FT) cells and SH-SY5Y neuroblastoma cells (**Fig. 1D)**. We then investigated the subcellular expression pattern with live imaging (**Fig. 1E,1F**). Live imaging frames of the basolateral imaging section showed that mTFP1 puncta localized to the same structures as mCherry as highlighted by line scan analysis (**Fig. 1E, 1F**). Furthermore, simultaneous two-channel live acquisition video shows colocalization of mTFP1 and mCherry-positive structures and their concomitant movement over time (**Fig. S2**; Supporting Movie File). Finally, we probed the lysates of FIRE-pHLy-expressing cells with an anti-LAMP1 antibody to confirm the size of the sensor between ∼130-160kD (**Fig. S3**). The two broad bands seen in the LAMP1 immunoblot suggest that the sensor is glycosylated, which was also seen in the sensor by Webb and colleagues^52^. Taken together, the microscopic and biochemical evaluation results confirm successful expression of the FIRE-pHLy cassette in HEK293FT and SH-SY5Y cells.

### FIRE-pHLy specifically localized to lysosomal compartments

We first investigated whether FIRE-pHLy expressed in HEK293FT cells sorted to lysosomal compartments. To do so, we tested colocalization of FIRE-pHLy with lysosomal, endosomal and mitochondrial sub-cellular markers (**Fig. 2A-E**). Cells were imaged using immunofluorescence confocal microscopy with three laser channels. Subsequently, we quantified the colocalization of FIRE-pHLy (using mCherry as reference) with existing markers for various sub-cellular organelles. We first assessed lysosomal markers by immunostaining for endogenous LAMP1 or LAMP2 or using LysoTracker Deep Red dye (Lyso-647). LAMP1 and LAMP2 are among the most abundant lysosome-associated membrane proteins ^68,69^. Endogenous LAMP1 and mCherry showed a strong positive correlation (r = 0.74 ± 0.03) (**Fig. 2A, 2F**). Similarly, LAMP2, a well characterized regulator of autophagy ^70^, colocalized with mCherry (r = 0.67 ± 0.04) (**Fig. 2B, 2G**). Lyso-647 is a widely used commercially available fluorescent probe that preferentially accumulates in acidic vesicular compartments, such as late endosomes and lysosomes ^71^. Co-localization of Lyso-647 and mCherry was similar to LAMP2 (r = 0.63 ± 0.03) (**Fig. 2C, 2H**). On the contrary, early endosome antigen 1 (EEA1) is a membrane-bound protein found specifically on early endosomes ^72^ and its labeling is characterized by large distinct ring-like structures ^73^. In contrast to the lysosomal markers, a lower fraction of mCherry associated with EEA1 (r = 0.41 ± 0.02) (**Fig. 2D,2I**) likely reflecting the maturation of FIRE-pHLy through the highly dynamic endolysosomal continuum ^74^. Finally, MitoTracker Deep Red (Mito-647) was used to stain mitochondria as a negative control (**Fig. 2E,2J**). Most Mito-647 exhibited minimal colocalization with FIRE-pHLy (r = 0.26 ± 0.03). The small percentage of colocalization was anticipated because mitochondria-lysosome crosstalk is known to occur ^75,76^.

**Figure 2.**
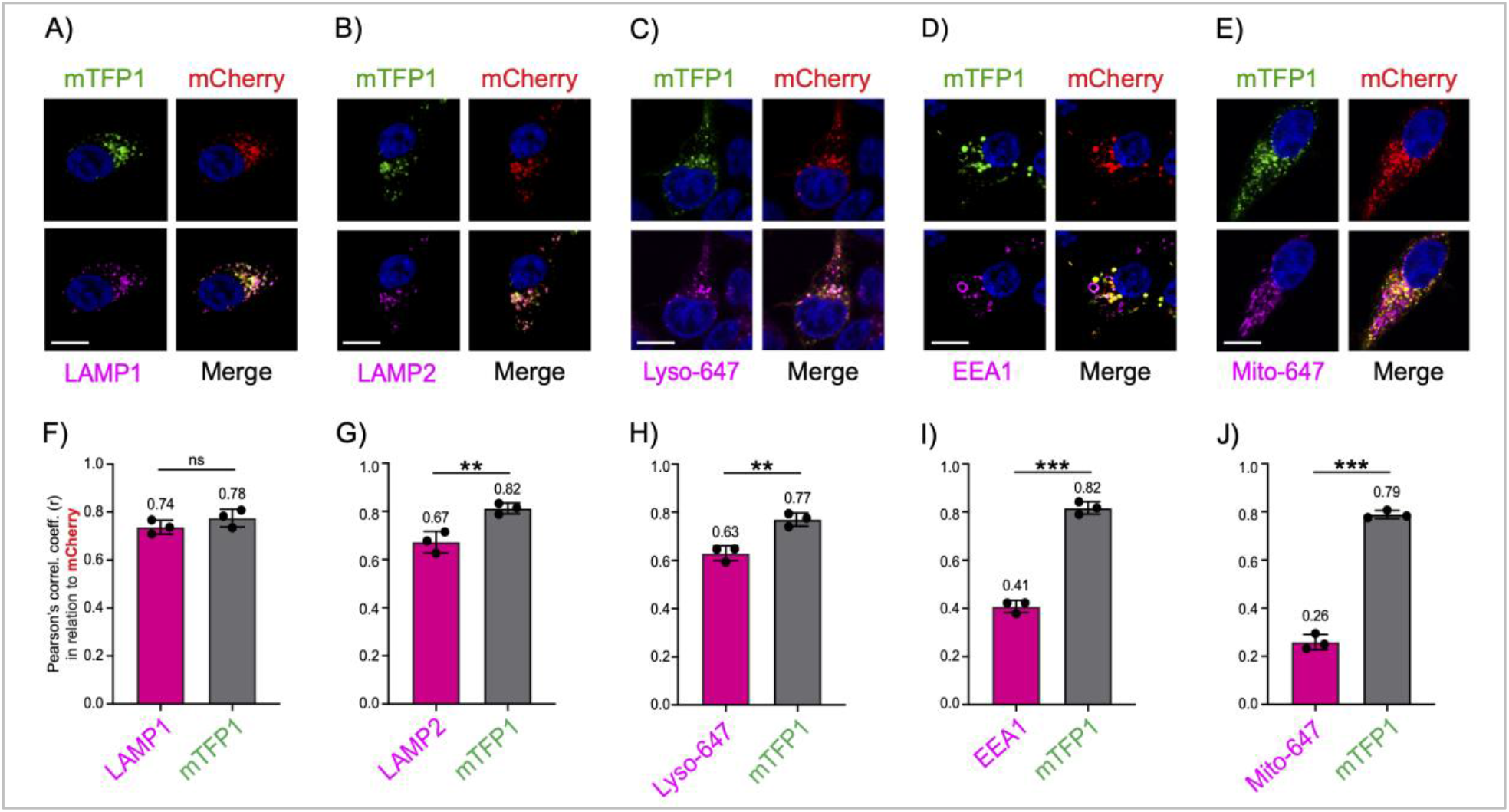
FIRE-pHLy localizes to lysosomal compartments. **(A-E)** Representative images of FIRE-pHLy-expressing HEK293FT cells stained with various markers (shown in magenta). **(A)** LAMP1 (lysosomal membranes), **(B)** LAMP2 (lysosomal membranes), **(C)** LysoTracker Deep Red or Lyso-647 (acidic compartments), **(D)** EEA1 (early endosomes), **(E)** MitoTracker Deep Red or Mito-647 (mitochondria). Nuclei are shown in blue. Scale bars = 10 μm. **(F-J)** Pearson’s correlation coefficients (r) calculated using the ImageJ plugin JACoP (Just Another Colocalization Plugin). Each graph shows a different marker colocalized with mCherry (magenta bars) and mTFP1 colocalized with mCherry (gray bars). Data points represents mean ± S.D. (3 independent replicates; n = 15 cells/replicate. Statistical analysis was performed using two-tailed, unpaired Student’s t-test. **p ≤ 0.01; ***p ≤ 0.001; ns = not significant).

As expected, since they are co-expressed as the same fusion protein, mTFP1 and mCherry showed consistently strong positive Pearson’s correlation coefficient values (within the range r = 0.78 – 0.82) across all images (**Fig. 2F-J**, gray bars). These coefficient values are less than 1.0 possibly due to mTFP1 quenching at physiological pH in lysosomes. Taken together, we concluded that mTFP1 and mCherry highly colocalizes with each other, as well as that FIRE-pHLy traffics through the endolysosomal sorting pathways to localize predominantly in lysosomal membranes.

### Quantification & visualization of pH-dependent, mTFP1 fluorescence in live cells

After confirming correct localization of FIRE-pHLy, we sought to demonstrate its pH sensitivity. Measuring intracellular and intralumenal pH of lysosomes using the ionophores, nigericin and monensin, is well established in previous protocols ^28,51,52,56,77–79^ and is currently the standard in the field (**Fig. 3A**). Nigericin (K^+^/H^+^) and monensin (Na^+^/H^+^) exchange K^+^ (and to a lesser extent Na^+^) for H^+^ across cell membranes, thus equilibrating external pH with that of the lysosomal lumen ^28,78^. Adapting these methods, we first used glass bottom chamber slides to qualitatively assess mTFP1 and mCherry fluorescence (using standard 488/561 nm filter sets) changes in HEK293FT cells at the applied pH values from 3.0 to 7.0. The fluorescence of mTFP1 increased from pH 3.0 to 7.0 while mCherry fluorescence remained relatively stable (**Fig. 3B**).

**Figure 3.**
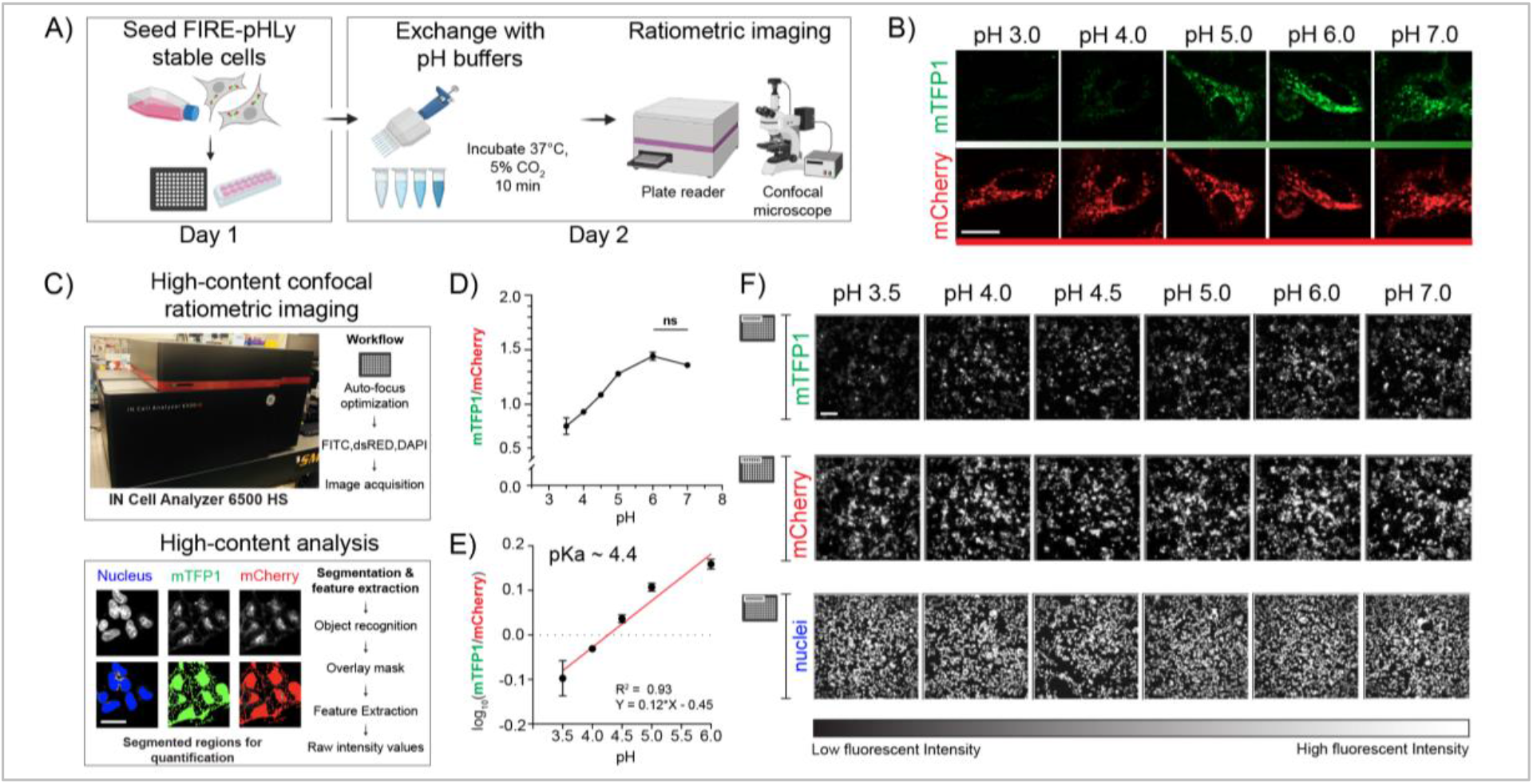
FIRE-pHLy biosensor responds to pH changes and is quantifiable with high-content analysis. **(A)** Workflow for pH calibration protocol. FIRE-pHLy-expressing cells were seeded into assay wells. Media was exchanged with pH buffers (at indicated values) supplemented with 10 μM nigericin and 1X monensin and was allowed to incubate for 10 min. Cells can be imaged live on either a confocal microscope or high-content plate reader. **(B)** Representative individual channel images of FIRE-pHLy-expressing HEK293FT cells imaged live by spinning disk confocal microscopy. Scale bar = 20 μm. **(C)** High-content analysis to quantify FIRE-pHLy fluorescence. Images were acquired on a plate-based confocal imager and analyzed on a custom-built segmentation protocol (see Methods). Masks for nucleus and FIRE-pHLy fluorescence were created and average mTFP1/mCherry ratios were calculated. **(D)** Cells were analyzed according to **(C)** and mTFP1/mCherry ratios were plotted against pH. Data points are presented as mean ± S.D., from 4 independent replicates; n = ∼10,000 cells quantified per pH value. Tukey’s test for multiple stepwise comparisons indicated significance between all pH groups, except 6.0 and 7.0. **(E)** Log_10_(mTFP1/mCherry) values between pH 3.5 and 6.0 were fit to a linear equation (R^2^ = 0.93). The pKa of FIRE-pHLy (in cells) was calculated to be ∼4.4. **(F)** Grayscale images of mTFP1, mCherry and nuclei taken from one random field of one representative assay well (of 96 well plate) at indicated pH values. Scale bar = 50 μm.

In order to increase the precision of measuring pH in a larger cell population, we adapted the assay to a high-content plate-based format. We built a lysosomal segmentation protocol (see Methods) that extracted fluorescence intensities of mTFP1 and mCherry, as well as nucleus count (**Fig. 3C**). From this analysis, we captured data from over 10,000 cells across four independent replicates at the applied pH values of 3.5 to 7.0 in 96-well plates (**Fig. 3C-F**) (see Methods). To quantify lysosomal pH, fluorescence intensity ratios for mTFP1 and mCherry (mTFP1/mCherry) were calculated and plotted according to the pH of the buffer. The ratio curve exhibited a significant positive relationship with pH, showing a ∼1.7-fold change in fluorescence ratio between pH 3.5 and 6.0 (**Fig. 3D**). Additional data indicates that mTFP1 fluorescence was the sole driver of the pH-dependent FIRE-pHLy ratio change (**Fig. S4**). Log_10_ transformation of ratios is linear from pH 3.5 to 6.0 (R^2^ = 0.93) (**Fig. 3E**).

It is noteworthy that though commonly used, the nigericin method has limitations. Equilibrating pH across membranes may affect the fluorescence intensity of both fluorophores. This sets a lower bound for pH calibration because the mCherry fluorophore is exposed to low pH. Furthermore, this method assumes that the applied pH represents the same pH to which mTFP1 was exposed. To validate the environment of mTFP1, we calculated the pKa of our ratiometric sensor to be ∼4.4 using a modified Henderson-Hasselbalch equation (Hoffmann & Kosegarten, 1995). This was in concordance with the *in vitro* mTFP1 pKa of ∼4.3 ^63^, suggesting that the pH of the lysosome was very similar to the pH of the applied buffer. Given the calibration challenges at low pH, we can establish that the fluorescence of FIRE-pHLy is sensitive to the applied pH in the range of 3.5 to 6.0; this range is appropriate for measuring pH in lysosomes under physiological conditions. Taken together, FIRE-pHLy fluorescence correlates with lumenal pH values in lysosomes.

### Functional validation of FIRE-pHLy in different cell types

Next, we evaluated the ability of FIRE-pHLy to monitor lysosomal pH under physiological conditions and pharmacological perturbations in widely used neurodegenerative disease cell models. We quantified the alkalinizing response to bafilomycin A1 (BafA1), a specific V-ATPase inhibitor, which functions by binding to the V0c subunit, thus blocking proton translocation ^81^. To select an appropriate BafA1 dose, we first tested multiple doses (30 nM to 1000 nM) in FIRE-pHLy-expressing HEK293FT cells and compared the sensor fluorescence to a pH calibration curve (**Fig. 4A-C**). To enable comparisons between samples (or potential high-throughput drug screening applications), we adapted the pH calibration protocol to fixed cells. Fixation led to 33.3% reduction of mTFP1 fluorescence and 10.6% reduction of mCherry fluorescence (**Fig. S5**), but did not change the overall ability to sense pH in the range of to 6.0. For this experiment, the calibration dynamic range became tighter showing a 1.59-fold change instead of 1.7-fold (**Fig. 3D**). Lysosomal pH increased dose-dependently with BafA1 334 concentration, plateauing at 300 nM with a pH∼5.6 compared to the control-treatment group pH of ∼4.1. Analysis of individual mTFP1 and mCherry fluorescence intensities under BafA1 treatment, confirmed that only mTFP1 fluorescence varies with lysosomal pH change (**Fig. S6**). A similar alkalinizing trend was observed in HEK293FT cells treated for 6 hours with 0.5 μM concanamycin A, another specific V-ATPase inhibitor ^82^ and with 30 μM chloroquine, a lysosomotropic drug known to inhibit autophagy and enlarge lysosomes ^83^ (**Fig. S7**).

**Figure 4.**
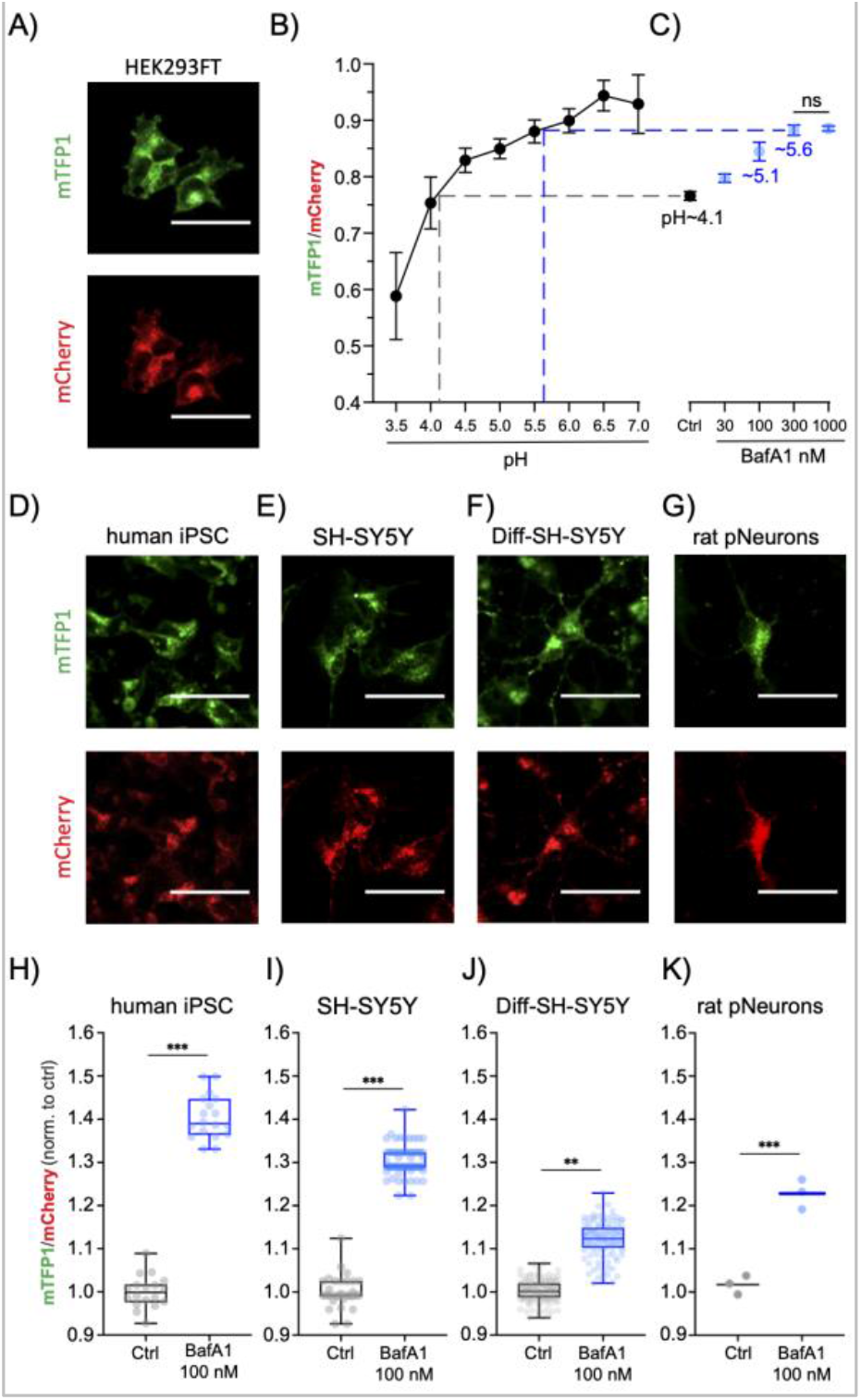
*In vitro* FIRE-pHLy models and relative pH measurements with bafilomycin A1. **(A)** Ratiometric images of 2% PFA-fixed FIRE-pHLy-expressing HEK293FT cells taken on a high-content imaging system (described in **Fig. 3**). **(B)** pH calibration curve generated from cells incubated with pH buffer (pH 3.5 – 7.0) and fixed with 2% PFA post 10min of treatment. Data points are presented as mean ± S.D., from 4 independent replicates; n = 10,000 quantified cells per pH value. **(C)** mTFP1/mCherry ratios of FIRE-pHLy-expressing HEK293FT cells treated with bafilomycin (BafA1 30 nM to 1000 nM) and 0.1% DMSO solvent control (Ctrl) for 6 hours prior to fixation and imaging. Data points are presented as mean ± S.D., from 6 independent replicates; n = 10,000 quantified cells per condition. Tukey’s test for multiple stepwise comparisons indicated significance between all groups including control, except BafA1 300 nM and 1000 nM. **(D-G)** Individual channel images (left to right) of FIRE-pHLy stably expressed in human iPSCs, SH-SY5Y, differentiated SH-SY5Y, and late embryonic rat hippocampal neuronal cells. All cells were fixed with 2%PFA prior to image acquisition. **(H-K)** 100 nM bafilomycin A1 was treated on cells for 6 hours and compared to 0.1% DMSO. Box-and-whisker plots show median, interquartile range (25th-75th percentile) and maximum/minimum values of mean ratios per well. **(H)** human iPSC; 18 independent wells in 96-well format; n = ∼15,000 quantified cells per well. 3 biological replicates. **(I)** SH-SY5Y; 76 independent wells in 384-well format; n = 2,500 cells per well. 2 biological replicates **(J)** RA-differentiated SH-SY5Y; 120 independent wells; n = 5,000 quantified cells per well. 4 biological replicates. **(K)** primary rat hippocampal neurons; 3 independent wells; n = 6,500 quantified cells per well. 1 biological replicate. Statistical analysis was performed using two-tailed, unpaired Student’s t-test. **p ≤ 0.01; ***p ≤ 0.001; ns = not significant. All scale bars = 25 μm.

Having established 100 nM as an appropriate BafA1 dose, we then probed for pH changes in induced pluripotent stem cells (iPSCs), SH-SY5Y neuroblastoma cells, retinoic acid-differentiated SH-SY5Y neuron-like cells and primary rat neurons (**Fig. 4D-G**), which were generated using lentiviral transduction of FIRE-pHLy. For transduction into these cells, the CMV promoter was exchanged for an UbC promoter-driven lentiviral FIRE-pHLy construct, since CMV is silenced by DNA methylation during differentiation and shows weak activity in certain cell types including iPSCs ^84,85^. Cells were treated with 100 nM BafA1 for 6 hours, fixed and subjected to high content analysis. Comparisons of mTFP1/mCherry fluorescence ratios with and without BafA1 treatment confirmed that FIRE-pHLy detected lysosomal alkalization across all cell lines tested (**Fig. 4H-K)**. The iPSCs had the largest ratio change of ∼40.4 ± 1.4% compared to control. On the other hand, differentiated SH-SY5Y cells had the smallest change. Though the change in ratio was only ∼11.9 ± 0.43%, using high-content analysis of over 5,000 cells, this change was statistically significant (p ≤ 0.01). Potential explanations for the observed cell type differences in the extent of relative pH response include differential BafA1 sensitivity or basal pH set point. For example, the expression and activity of V-ATPases is regulated differentially in mammalian cells ^86^. Cell type-dependent pH regulatory and compensatory mechanisms warrant further investigation.

Overall, our data demonstrates that FIRE-pHLy can be targeted to lysosomes in multiple neurodegenerative disease cell models. This opens future avenues to profile lysosomal pH dynamics in cellular systems harboring different genetic mutations and further use for applications in lysosome-based drug discovery.

## CONCLUSIONS

In summary, we have developed FIRE-pHLy, a genetically encoded ratiometric pH biosensor that localizes to lysosomal membranes and measures lumenal pH within physiological ranges (3.5 to 6.0). FIRE-pHLy responds robustly to pH changes and is amenable to stable integration to multiple cellular models, including differentiated and primary cells. Moreover, FIRE-pHLy is amenable to live- and fixed-cell assays, as well as both high-resolution confocal microscopy and quantitative high-content imaging. We anticipate that FIRE-pHLy will be applied to elucidate pH dynamics in basic lysosomal biology and disease. Moreover, the ability to quantifiy the sensor in 96-well plates with high-content analysis enables its translation to phenotypic-screening platforms for drug discovery in fields such as neuroscience, immunology and cancer biology. Finally, this study opens new avenues to profile lysosomal functions in animal models of childhood or age-associated neurological diseases.

## METHODS

### Construction of <u>F</u>luorescence <u>I</u>ndicator <u>RE</u>porting <u>pH</u> in <u>Ly</u>sosomes (FIRE-pHLy)

The genetically encoded FIRE-pHLy reporter cassette consists of the following coding segments, from the N-terminus: CMV-human LAMP1 signal peptide (84bp) – mTFP1 – flexible linker 1 (GGSGGGSGSGGGSG) – human LAMP1 – rigid linker 2 (PAPAPAP) - mCherry. Source of different elements are as follows: LAMP1 signal peptide and human LAMP1 were PCR amplified from LAMP1-mGFP (Addgene Plasmid #34831, kind gift from the Mark Von Zastrov lab, University of California, San Francisco, UCSF), mTFP1 amplified from mTFP1-pBAD (Addgene Plasmid #54553), mCherry amplified from pcDNA3.1-mCherry (Addgene Plasmid #128744). The DNA segments were PCR amplified (Phusion High-Fidelity PCR Master Mix, NEB, UK, #M0531) and fused with Gibson recombination cloning method (Gibson Assembly Master Mix, NEB, UK, #E2611) in pEGFP-N3 empty backbone. The linker sequences were incorporated into the primer sequences. The FIRE-pHLy expression cassette was cloned into lentiviral vectors with CMV promoter (pLJM1-EGFP; Addgene Plasmid #19319) and hUbC promoter (FUGW; Addgene Plasmid #14883) by Epoch Life Science services (Sugar Land, Texas, USA).

### Cell culture and lentiviral transduction

All cells were cultured at 37°C with 5% CO_2_ atmosphere and maintained under standard procedures. HEK293FT cells (Thermo Fisher Scientific; Carlsbad, CA, USA, #R70007) were cultured in Dulbecco’s Modified Eagle medium (DMEM; Life Technologies, Carlsbad, CA, USA, #11-995-073) with 10% heat-inactivated fetal bovine serum (FBS; Gemini Bio, Sacramento, CA, USA, #GEMZR039) containing 1% penicillin and streptomycin (pen/strep) (Gibco; Thermo Fisher Scientific, Inc., Waltham, MA, USA, #15140122) with 500 μg/mL G418 sulfate antibiotic (Thermo Fisher Scientific, #11811031). SH-SY5Y cells (American Type Culture Collection; ATCC, Maryland, USA, #CRL-2266) were maintained in 1:1 Eagle’s Minimum Essential Medium (EMEM; ATCC, #30-2003) and F12 medium (Life Technologies; Carlsbad, CA, USA, #11765062) with 10% FBS and 1% pen/strep. Cells were trypsinized with 0.05% Trysin-EDTA solution (Sigma, St. Louis, MO, USA, #T4049) during routine passaging. Lentivirus production and titer assessments of FIRE-pHLy-lentivirus were performed by the UCSF ViraCore facility. For lentivirus transduction, HEK293FT and SH-SY5Y cells were plated in 6-well plates and cultured to ∼ 70% confluence. Protocol was modified for iPSCs and primary rat neurons (see below). Lentivirus infections were carried out in the presence of 10 μg/mL of polybrene (Sigma, St. Louis, MO, USA, #S2667) in complete media. 48 hours post-transduction, cells were selected with 1 μg/mL puromycin (Millipore; Carlsbad, CA, USA, #540411) to generate stable transgene-expressing cell lines. Long-term transgene expression was maintained by selecting for resistance to puromycin at a final concentration of 1 μg/mL. SH-SY5Y cells were sorted for green and red positive fluorescence signal on a SH800S Cell Sorter (Sony Biotechnology) at the UCSF Laboratory of Cell Analysis.

### Generation of FIRE-pHLy-expressing iPSCs

The F11350 iPSC line was obtained from the laboratory of Celeste Karch at Washington University School of Medicine ^87^. Cells were maintained in matrigel (Corning, #354277) coated plates using mTSER media (StemCell Technologies, Vancouver, Canada, #05850), which was replaced every day. During passaging, cells were lifted using Accutase (ThermoFisher Scientific, #A1110501) and then replated in media supplemented with 10 µM ROCK inhibitor (Y-27632, StemCell Technologies, #72304). For the transduction of the virus, iPSCs were plated onto matrigel-coated 24-well plates at a density of 50,000 cells per well. Serial dilutions of the UbC promoter FIRE-pHLy lentiviral vector were prepared in mTSER media with 4 µg/ml polybrene (Sigma, #S2667) Lentivirus media was allowed to transduce cells for 24 hours and then fresh media changes were performed every day until 80% confluence was reached. Clonal populations of green/red fluorescence positive cells were manually selected and transferred into separate wells for expansion.

### Isolation of primary rat neurons and FIRE-pHLy lentivirus transduction

Wildtype SAS Sprague Dawley rats (Charles River Laboratories, Wilmington, MA, USA) and isolation reagents were a kind gift from the Molofsky Lab (UCSF). Embryos staged at day 18 were dissected from one pregnant rat and immediately placed in chilled brain dissection buffer (HBSS-Ca^2+^/Mg^2+^-free with 10 mM HEPES buffer, pH 7.3). Brains were removed from 5 individual embryos and hippocampi halves were isolated after removal of the meninges. Hippocampi were digested with trypsin/EDTA solution and DNAase at 37°C incubation for 25 min. Quenching buffer (HI-OVO diluted 1:5 in HBSS-Ca^2+^/Mg^2+^-free, with 50% glucose, ovomucoid, bovine serum albumin, and DNAase) was subsequently added to inhibit trypsin digestion. Following centrifugation and buffer removal, culture medium I (DMEM-high glucose, L-glutamine-sodium pyruvate free) supplemented with 10% fetal calf serum (FCS) (not heat activated) was added to partially digested hippocampi. Cells were then manually dissociated and plated in poly-D-lysine coated 96-well plates (Greiner Bio-One, Kremsmünster, Austria, #655956) at a density of 15,000 cells per well. After 24 hours, media was replaced with culture medium II (Neurobasal medium, 1% heat inactivated FCS, 2% B27 supplement, 1X Glutamax I, 1X MycoZap plus, and 15mM NaCl). At DIV 5 (5 days in vitro), 5 μM 5-Fluorouracil was added to curb glial cell proliferation. At DIV 7, neurons were transduced with UbC-FIRE-pHLy lentivirus for 24 hours. Half media changes were performed every two days until DIV 14.

### RA-differentiation of FIRE-pHLy SH-SY5Y neuroblastoma cells

SH-SY5Y cells were seeded in collagen type I-coated μClear 96-well plates (Greiner Bio-One, #655956) at a density of 10,000 cells/cm^2^ 24 hours before start of differentiation. Differentiation media was composed of 10 µM of retinoic acid (RA) (Sigma, #R2625) in EMEM/F12 media supplemented with 10% FBS and 1% P/S. After 6 days of RA treatment, cells were treated for 4 days with 50 ng/mL of brain-derived growth factor (BDNF) (Peprotech, Rocky Hill, NJ, USA, #450-02B) in serum-depleted EMEM/F12 media supplemented with 1% pen/strep ^88,89^.

### Antibodies & reagents

#### Immunofluorescence

AlexaFluor 647 mouse-anti-hLAMP1 (1:500, Biolegend, Carlsbad, CA, USA; #328611), AlexaFluor 647 mouse-anti-hLAMP2 (1:500, Biolegend, Carlsbad, CA, USA; #354311), mouse-anti-EEA1 (1:1000, BD Biosciences; Franklin Lakes, NJ, USA; #610457), AlexaFluor 647 goat anti-mouse (1:500, Life Technologies; #A21206). *Western blot*. hLAMP1 (1:1000, DSHB, University of Iowa, USA, #2296838). *Reagents*. LysoTracker Deep Red (Life Technologies; #L12492), MitoTracker Deep Red FM (Invitrogen; Carlsbad, CA, USA, #M22426), Monensin solution 1000X (Invitrogen; #501129057), Nigericin solution (Sigma Aldrich, USA, #SML1779), paraformaldehyde (PFA) solution (Fisher Scientific; #50980494), glycine (Sigma; #G7126), bovine serum albumin (BSA) (Fisher Scientific; #BP1605100), D-PBS (Sigma; #D8662), Saponin (Sigma; #S7900).

### Western blotting

Lysates from FIRE-pHLy-expressing HEK293FT cells were collected from 1X RIPA buffer (Fisher Scientific, #89900) supplemented with a cocktail of phosphatase and protease inhibitors (Roche, Basel, Switzerland, #4693116001) and 1 ug/mL peps tatin A (Thermo Scientific, #78436). Sample protein concentrations were determined using the Pierce BCA Protein Assay kit (Thermo Scientific, #PI23225). Samples were loaded onto a Novex NuPAGE SDS-PAGE gel system with MOPS running buffer (Life Technologies, #NP001). Proteins were transferred onto nitrocellulose membranes and blotted with indicated antibodies. Imaging of band intensities were performed on a LI-COR Odyssey Infrared System (LI-COR Biosciences, Lincoln, NE, USA).

### Live imaging, Immunofluorescence microscopy and colocalization analysis

FIRE-pHLy-expressing HEK293FT cells were plated in TC-grade chamber slides (μ-Slide 8-well chamber slide, Ibidi, Gräfelfing, Germany, #50305795) at a density of 30,000 cells per well for 24 hours. For live uptake of organelle markers, cells were incubated with 30 nM LysoTracker Deep Red or 30 nM MitoTracker Deep Red FM along with 1:1000 Hoechst dye (10mg/mL Hoescht 33342 solution, ThermoFisher, #H3570) in culture medium at 37°C/5% CO_2_ for 10 min. For live microscopy, FIRE-pHLy-expressing HEK293FT stable cells were grown on μ-Slide 8-well chamber slide (Ibidi, Gräfelfing, Germany, #50305795) with culture medium supplemented with 10mM HEPES. Time-lapse imaging was performed at 37 °C using a spinning-disk microscope, NikonTi (Inverted), UCSF Facility through a Plan Apo VC 100x/1.4 Oil objective lens. The apparatus is composed of a Andor Borealis CSU-W1 spinning disk confocal, Andor 4-line laser launch (100 mW at 405, 561, and 640 nm; 150 mW at 488 nm), equipped with an Andor Zyla sCMOS camera (5.5 megapixels) for image acquisition, and Micro-Manager 2.0 beta 3 software to control the setup. The images were acquired simultaneously with configuration parameters (100ms exposure) GFP and mCherry channels with 25% laser power. For immunofluorescence staining (LAMP1, LAMP2, and EEA1), cells were washed once with 1X D-PBS (with MgCl_2_ and CaCl_2_) and fixed with 2% PFA for 15 min at room temperature (RT). Then, cells were washed once with glycine, blocked for 2 min with 2% BSA/D-PBS, and permeabilized with 0.01% saponin/2% BSA for 1 min at RT. Cells were incubated with primary antibodies for 1 hour at RT, secondary antibodies for 1 hour at RT shielded from light, and washed twice. Cells were imaged using an inverted confocal line-scanning microscope (DMi8 CS Bino, Leica Microsystems Inc., Wetzlar, Germany) with a 63x/1.40 oil-immersion objective lens at 2048×2048 pixel resolution. Fluorescence images was acquired with sequential scanning between frames on the LAS X SP8 Control Software system using preset channel settings (Blue Ex/Em: 405 nm/410-464nm; Green Ex/Em: 470 nm/474-624nm; Red Ex/Em: 587 nm/592-646 nm; Far Red Ex/Em: 653 nm/658-775 nm). Randomly imaged fields were processed (background subtraction, thresholding) and the cytosolic green/red values with Pearson’s correlation coefficients were calculated using ImageJ (NIH, Maryland, USA) ^90^ plugin, JACoP (Just Another Co-localization Plugin) ^91^ and linescan analysis performed using ImageJ (NIH, Maryland, USA) ^90^.

### pH calibration buffers, generation of standard curve and pKa calculation

pH calibration buffers and procedures were adapted from a previously described study with few modifications ^28^. Buffer recipe is described below, composed of 5 mM NaCl, 115 mM KCl, and 1.3 mM MgSO_4_•7H_2_O, 25 mM MES buffer with pH adjusted in the range 2.0 – 7.0. Freshly made buffers were supplemented with 10µM nigericin and 1X monensin.

| Stock Conc. | 5M NaCl | 1M KCl | 1M MgSO<sub>4</sub>•7H<sub>2</sub>O | 0.5M MES | 6.7mM Nigericin | Monensin (1000X) | 1N KOH/<br>1N HCl * | Final volume |
| --- | --- | --- | --- | --- | --- | --- | --- | --- |
| Volume | 10 μl | 1150 μl | 13 μl | 500 μl | 15 μl | 10 μl | * | 10ml |
| Final Conc. | 5 mM | 115 mM | 1.3 mM | 25 mM | 10 μM | 1X |  | (H<sub>2</sub>O) |

\*pH values are adjusted using pH meter with appropriate acid/base. (conc., concentration)

On day 1, FIRE-pHLy-expressing HEK293FT cells were seeded at a density of 10,000 cells/well resuspended in 100 μl media in collagen type-1 coated, black-bottom 96-well plates (µClear, Greiner Bio-One; #655956). Plates were left in the culture hood at room temperature for 45 min to allow for even cell distribution before incubated at 37°C/5.0% CO_2_ overnight. On day 2, cell nuclei were stained with 1:1000 (vol/well) Hoechst dye (10 mg/mL Hoescht 33342 solution, ThermoFisher, #H3570) diluted in cell culture media for 20 min at 37°C/5.0% CO_2_. After one wash with 50 μl of 1X D-PBS, 50 μl of each pH titration buffer supplemented with 10 µM nigericin and 1X monensin were added to wells and incubated at 37°C/5.0% CO_2_ for 10 min. Note: to attain uniform exposure to pH buffers including ionophores, samples should be (i) imaged within 10-15 min after buffer addition ^28,56,77,92^, (ii) fixed after 10-15 min, or (iii) manually imaged one at a time. Plates were then immediately imaged (total imaging time > 5 min) live on the IN Cell Analyzer 6500 HS (General Electric Life Sciences/Cytiva, Marlborough, MA, USA) and processed (see *High-content confocal microscopy, feature extraction and ratiometric image analysis)*. Liquid dispensing and aspiration were performed using an automatic multichannel pipette (Voyager II, INTEGRA Biosciences Corp, Hudson, NH, USA; #4722). After raw mTFP1 and mCherry intensity values were obtained, log10(mTFP1/mCherry) values were fit with a linear regression. A modified Henderson-Hasselbalch equation, as previously used ^80^, was used to calculate pKa.

### Lysosomal inhibitor assay in cells

Cells were seeded and cultured on 96-well plates prior to addition of inhibitors - 100 nM bafilomycin A1 (LC Laboratories, Woburn, MA #B-1080), 30 μM chloroquine (Sigma, #C6628), and 0.5 μM concanamycin A (Sigma, #C9705). After 6 hours, cells were fixed with 2% PFA at room temperature (RT) for 15 min and washed once with 1X D-PBS. Cells were stained with 1X Hoechst dye for 20 min at RT protected from light and washed once with 1X D-PBS. Plates were imaged on the IN Cell Analyzer 6500 HS and processed (see *High-content confocal microscopy, feature extraction and ratiometric image analysis)*.

### High-content confocal microscopy, feature extraction and ratiometric image analysis

Black 96-well assay plates (µClear bottom, Greiner Bio-One; #655956) were imaged using a fully automated laser-scanning confocal cell imaging system (IN Cell Analyzer 6500HS, GE Life Sciences) with a NIKON 20X/0.75, Plan Apo, CFI/60 objective lens and preset excitation lasers (Blue 405 nm; Green: 488 nm; Red: 561 nm) with simultaneous acquisition setting. Laser and software autofocus settings were applied to determine a single optimal focus position. The EDGE confocal setting was used to increase image resolution and improve downstream visualization and segmentation of lysosomes. Nine images were acquired per well and were distributed in a 3×3 equidistant grid positioned in the well center. Wells were imaged sequentially in a vertical orientation. Image stack files were analyzed on a high-content image analysis software (IN Cell Developer Toolbox v1.9, GE Life Sciences). Target set segmentation and quantification measures were developed for individual channels and applied onto all sample images. Cell nuclei were segmented using a preset nucleus type segmentation module. The size of nuclear mask was adjusted according to the cell type. Visual inspection of several reference fields across multiple wells confirmed segmentation accuracy. Total number of segmented nuclei was quantified per well. To quantify FIRE-pHLy fluorescence, mCherry was used as the reference channel for segmenting lysosomes. mCherry fluorescence provided a robust representation of FIRE-pHLy localization for the purposes of delineating lysosomal objects, compared to mTFP1 fluorescence, whose signal varies with lysosomal pH. A preset vesicle segmentation module was applied on the 561nm source images with acceptance criteria (Dens-levels>300), min/max granule size (1-10um), scales = 2, sensitivity = 33, low background, and no shape constraints settings. These settings created an object “mask” for lysosomes, which was directly applied to 488nm source images to segment mTFP1 identically as mCherry. Mean fluorescence intensities of mCherry and mTFP1 channels were generated and the ratios calculated. All measures were outputted as a Microsoft Excel file for further analysis.

### Data presentation, statistical analysis, and illustrations

All data were generated from randomly selected sample populations from at least three independent experiments represented unless otherwise mentioned in corresponding figure legends. Statistical data were either presented in box–and-whisker plots with median, interquartile range, and maximum & minimum values or bar graphs with mean ± S.E.M. Multiple comparisons between groups were analyzed by one-way ANOVA with Tukey’s test and statistical significance for two sets of data was determined by two-tailed, unpaired Student’s t-test. All data plots and statistical analyses were performed using GraphPad Prism 8 with no samples excluded. Significant differences between experimental groups were indicated as *P<0.05; **P<0.01; ***P<0.001; Only P <0.05 was considered as statistically significant. NS = not significant. Pre-processing of data was organized in Microsoft Excel. Cartoon schematics were created on Biorender.com. Figures were assembled on Adobe Illustrator.

## DATA AND REAGENT AVAILABLE

The authors declare that all relevant data supporting the findings of this study are available within the paper. Any data or reagents can be obtained from the corresponding authors (A.W.K. and M.R.A.) on reasonable request.

## SUPPORTING INFORMATION AVAILABLE

The following files are available free of charge.

ACS Sensors_Chin et al_Supporting Information.pdf

ACS Sensors_Chin et al_Supporting Movie.mpeg

The supporting information shows cross-excitation images of FPs, time-lapse movie of mTFP1 and mCherry-positive lysosomes in cells, immunoblot of FIRE-pHLy-expressing cells, analyses of individual mTFP1 and mCherry fluorescence under pH calibration, fixation, bafilomycin A1 treatment conditions, application of different lysosomal inhibitors to elevate pH in lysosomes.

## ACKNOWLEDGEMENTS

We thank the Mark Von Zastrov lab (UCSF) for providing us with the hLAMP1-GFP plasmid. We also acknowledge the UCSF Small Molecule Discovery Center for their tremendous support and advice on our high-content imaging and analysis protocols. Additionally, we thank members of the Kao and Arkin labs for their thoughtful discussions. This work has received support from the NIH/NIA (<u>R01 AG058447</u>).

## AUTHOR CONTRIBUTIONS

M.Y.C., A.R.P., M.G, M.R.A. and A.W.K for conceptualization and design; M.Y.C., A.R.P., K.A., A.L.W., C.A. and M.W. for experimental design and data acquisition; P.T.N. and A.V.M for conceptualization, design and data acquisition of primary rat neuron experiments; all authors analyzed and interpreted data; M.Y.C., A.R.P., M.R.A. and A.W.K. wrote the manuscript; all authors edited the manuscript.

## Abbreviations

FIRE-pHLy: <u>F</u>luorescence Indicator <u>RE</u>porting <u>pH</u> in <u>Ly</u>sosomes
LAMP1: Lysosomal-associated membrane protein 1
LAMP2: Lysosomal-associated membrane protein 2
V-ATPase: Vacuolar-type ATPase
FP: Fluorescent Protein
mTFP1: monomeric teal fluorescent protein
DAMP: (N-(3-((2,4-dinitrophenyl)amino)propyl)-N-(3-aminopropyl)methylamine, dihydrochloride
FITC: fluorescein isothiocyanate
BafA1: Bafilomycin A1
ConA: Concanamycin A
CQ: Chloroquine
iPSC: Induced pluripotent stem cells
HEK293: human embryonic kidney 293
RA: Retinoic acid
BDNF: Brain-derived neurotrophic factor
CMV: cytomegalovirus
UbC: ubiquitin C.

