## Supporting Information for "A genetically encoded, pH-sensitive mTFP1 biosensor for probing lysosomal pH"

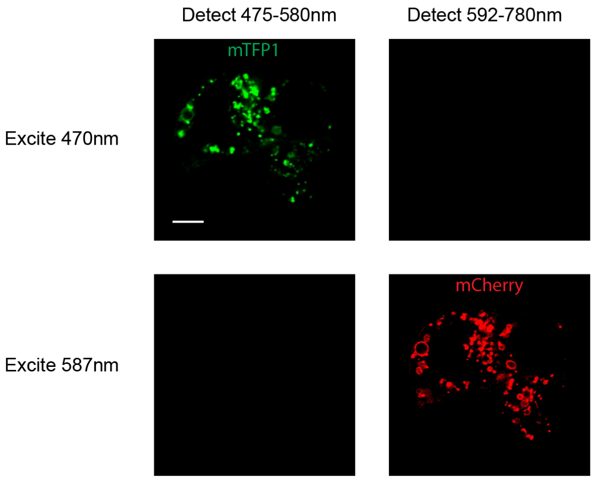


**Fig S1. Cross excitation of mTFP1 and mCherry.**
Confocal images of FIRE-pHLy-expressing HEK293FT cells. mTFP1 is excited at 470 nm and detected between 475-580 nm, but not between 592-780 nm. Conversely, mCherry is excited at 587 nm and detected between 592-780 nm, but not between 475-580 nm. Scale bar = 10 µm.

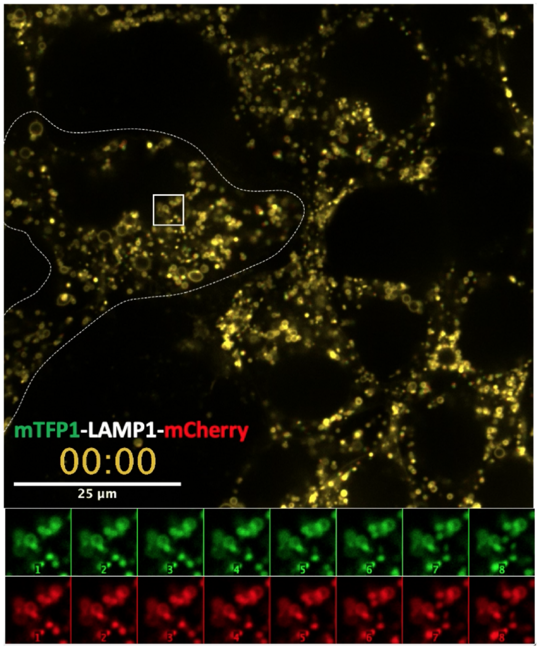


**Fig S2. Expression and live imaging of FIRE-pHly in HEK293FT stable cells.**
Time-lapse merged video (see Supporting Movie File) acquired using spinning-disc confocal microscopy on HEK293FT stable cells expressing chimeric construct encoding FIRE-pHLy. Note the consecutive time-lapse images (first 8 frames, lower panel) where mTFP1 (green) positive dynamic structures colocalize and show concomitant movements with mCherry (red) labeled LAMP1 positive lysosomes. Simultaneously acquisition using GFP/mCherry channel with 100 ms exposure, video was shown at 7 frames/s. Scale bar = 25 µm.

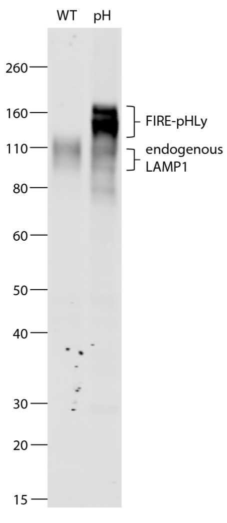


**Fig S3. Western blot analysis of FIRE-pHLy expression in HEK293FT cell lysates.**
Lysates from wild-type (WT) and FIRE-pHLy-expressing (pH) HEK293FT cells were immunoblotted with an anti-hLAMP1 antibody to detect pH sensor expression levels. The observed FIRE-pHLy molecular weight (MW) is ~130-160 kD. The observed MW of LAMP1 due to glycosylation is ~90-120 kD (note: calculated MW of LAMP1 is ~40 kD). The MWs mTFP1 and mCherry are 27 kD each. Protein loading: 20 ug/lane.

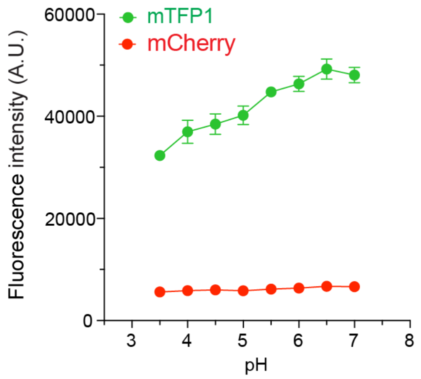


**Fig S4. Measured fluorescence intensities of FIRE-pHLy FPs in cells calibrated with pH buffers.** Individual mTFP1 and mCherry fluorescence intensities plotted against pH (3.5-7.0). FIRE-pHLy-expressing HEK293FT cells were incubated with pH buffers (3.5-7.0) supplemented with 10 µM nigericin and 1X monensin and imaged with a high-content plate reader. Data points are presented as mean ± S.D., from 6 independent wells; n = ~10,000 cells quantified per pH value.

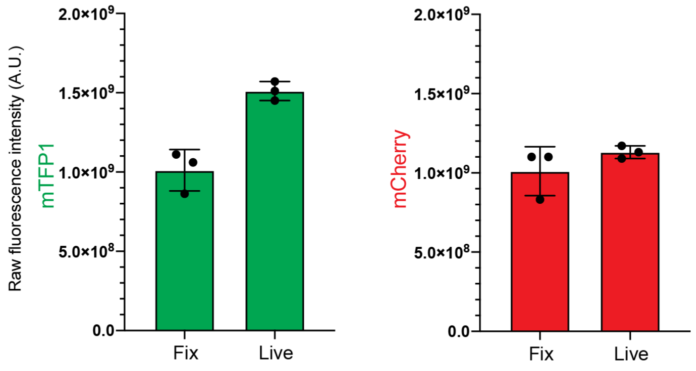


**Fig S5. Fixed- and live-cell fluorescence measurements for mTFP1 and mCherry FPs.**
Raw mTFP1 and mCherry fluorescence intensities measured from FIRE-pHLy-expressing HEK293FT cells that were either imaged live (in culture media, pH 7.4) or post-PFA fixation (in PBS, pH 7.4). Data points are presented as mean ± S.D., from 3 independent wells; n = ~5,000 quantified cells per well.

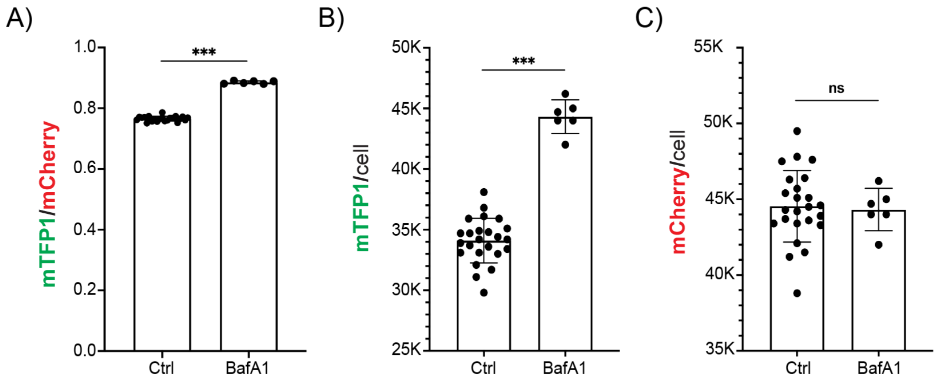


**Fig S6. Ratiometric validation of individual FIRE-pHLy fluorophores under BafA1 conditions.** (A) mTFP1/mCherry ratio quantified from FIRE-pHLy-expressing HEK293FT cells treated with 1 μM bafilomycin for 6 hours compared to 0.1% DMSO solvent control. (B) mTFP1 mean fluorescence intensity normalized by cell count. (C) mCherry mean fluorescence intensity normalized by cell count. Data points are presented as mean ± S.D., from 6 independent replicates; n=quantified 7,500 cells per replicate. Statistical analysis was performed using two-tailed, unpaired Welch's t-test for unequal variances. ***$p \leq 0.001$; ns = not significant.

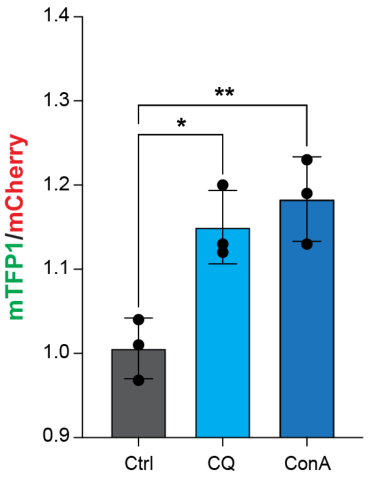


**Fig S7. pH elevation with lysosomal pharmacological inhibitors.**
Ratiometric measurements (mTFP1/mCherry) taken from FIRE-pHLy-expressing HEK293FT cells treated with 0.1% DMSO (Ctrl), 30 µM chloroquine (CQ) and 0.5 µM concanamycin A (ConA) for 6 hours before fixation. Data points are presented as mean ± S.D., from 3 independent wells; n = ~5,000 quantified cells per well. Statistics were conducted with one-way ANOVA for multiple comparisons. *$p \leq 0.05$; **$p \leq 0.01$.
